# Virtual Screening–Guided Discovery of Small Molecule CHI3L1 Inhibitors with Functional Activity in Glioblastoma Spheroids

**DOI:** 10.1101/2025.07.31.667816

**Authors:** Baljit Kaur, Katrin Denzinger, Longfei Zhang, Nelson García-Vázquez, Gerhard Wolber, Moustafa Gabr

## Abstract

Chitinase-3-like protein 1 (CHI3L1), a glycoprotein implicated in inflammation, fibrosis, and cancer, has emerged as a potential therapeutic target for glioblastoma (GBM). CHI3L1 contributes to tumor progression and immune evasion by promoting STAT3 signaling and mesenchymal transition. To identify small molecule CHI3L1 inhibitors, a structure-based 3D pharmacophore model was developed and applied to virtually screen over 4.4 million compounds from the Enamine collection. Following multi-tiered filtering, 35 candidates were selected for experimental evaluation. Binding validation via microscale thermophoresis (MST) confirmed dose-dependent CHI3L1 interactions for two compounds, **8** and **39**, with dissociation constants (K_d_) of 6.8 µM and 22 µM, respectively. These affinities were further supported by surface plasmon resonance (SPR), which yielded K_d_ values of 5.69 µM for compound **8** and 17.09 µM for compound **39**. In 3D GBM spheroid models, compound **8** significantly reduced spheroid viability and attenuated phospho-STAT3 levels, consistent with CHI3L1 pathway disruption. These findings identify two promising scaffolds and support the utility of pharmacophore-guided virtual screening for discovering functionally active ligands targeting CHI3L1 in GBM.

Table of Contents artwork

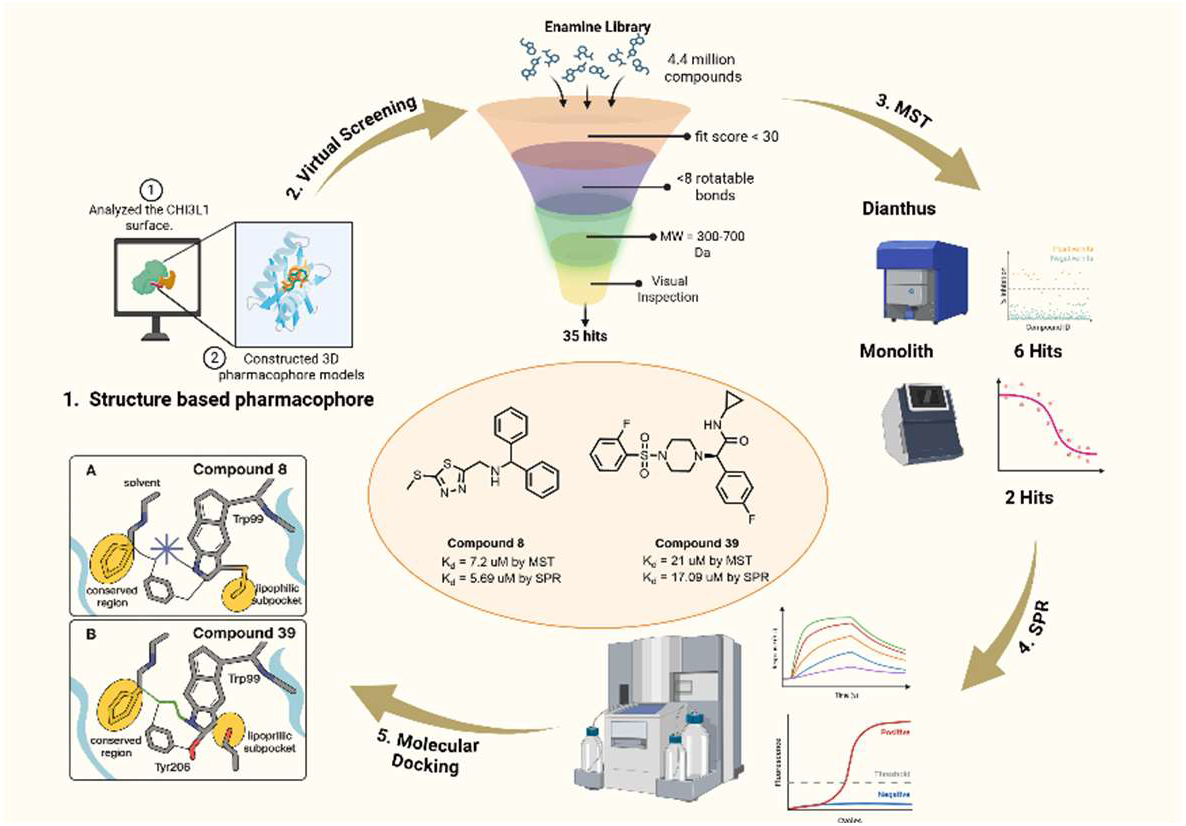

---

CHI3L1 (Chitinase-3-like protein 1), commonly referred to as YKL-40, is a secreted glycoprotein with diverse biological functions spanning inflammation,^1–4^ extracellular matrix remodeling, ^5–7^ angiogenesis,^8–10^ cell survival,^11,12^ and immune modulation. ^13,14^ Although it shares structural features with active chitinases, CHI3L1 lacks enzymatic activity due to specific amino acid substitutions in its catalytic domain.^15,16^ Rather than functioning as an enzyme, its biological activity is mediated through interaction with several cell surface receptors and binding partners.^17^ CHI3L1 is produced by a variety of cells, including macrophages, neutrophils, epithelial cells, fibroblasts, and tumor cells-especially in response to tissue injury, stress, or chronic inflammation.^18–28^

A defining characteristic of CHI3L1 is its marked upregulation in numerous pathological conditions. In chronic inflammatory disorders such as asthma,^29,30^ rheumatoid arthritis,^31^ and inflammatory bowel disease,^32^ CHI3L1 contributes to immune cell recruitment, perpetuates inflammation, and exacerbates tissue injury. In fibrotic diseases, including liver fibrosis,^33–35^ idiopathic pulmonary fibrosis,^30,36–38^ and systemic sclerosis,^18,39,40^ CHI3L1 promotes fibroblast activation and excessive deposition of extracellular matrix components, leading to progressive scarring and organ dysfunction. Within the tumor microenvironment, CHI3L1 supports malignant progression by enhancing cell proliferation, promoting angiogenesis, inhibiting apoptosis, and enabling immune escape. Its expression often correlates with advanced disease stages and unfavorable prognoses, underscoring its dual role as both a biomarker and an active driver of disease.

The biological activity of CHI3L1 is mediated through its interactions with multiple receptors and effector pathways that contribute to immune modulation, tissue remodeling, and tumor progression. CHI3L1 has been reported to bind several receptors, including IL-13 receptor alpha 2 (IL-13Rα2),^41^ syndecan-1,^42^ the receptor for advanced glycation end-products (RAGE),^43^ CD44,^44^ and various integrins.^45^ Additional studies have described binding to Galectin-3 (Gal-3),^46^ a β-galactoside-binding lectin implicated in a wide range of pathological processes.^47–50^

More recent findings have revealed a direct and functionally consequential interaction between CHI3L1 and signal transducer and activator of transcription 3 (STAT3), an established oncogenic driver in glioblastoma (GBM) and other solid tumors.^51^ STAT3 is activated by phosphorylation and translocates to the nucleus, where it drives the expression of genes that promote cell survival, proliferation, angiogenesis, and immune evasion.^52-55^ As a key regulator of tumor growth and therapy resistance, STAT3 is critically involved in GBM progression and represents a viable therapeutic target.^56-59^ In GBM, STAT3 has been shown to sustain a pro-tumorigenic transcriptional program that supports glioma stem cell maintenance, suppresses antitumor immunity, and facilitates proneural-to-mesenchymal transition (PMT), a process associated with therapy resistance and poor prognosis.^51^ CHI3L1 interacts with the coiled-coil domain of STAT3, stabilizing its active conformation and promoting nuclear translocation and transcriptional activity.^51^ This signaling axis directly upregulates the expression of SPP1 (osteopontin), a key mediator of mesenchymal reprogramming and glioma plasticity.^51^

In light of these findings, the CHI3L1-STAT3 pathway represents an emerging therapeutic vulnerability in GBM. Pharmacological inhibition of this axis using the small molecule hygromycin B (HB) has been shown to disrupt CHI3L1-STAT3 binding, reduce STAT3 phosphorylation and nuclear translocation, and attenuate SPP1 expression.^51^ In preclinical models, HB treatment suppressed tumor growth, reversed mesenchymal features, and reprogrammed the immune microenvironment.^51^ In this context, our study aims to identify small molecule CHI3L1 binders that disrupt its functionally relevant binding cleft, thereby interfering with downstream STAT3 activation and transcriptional reprogramming in GBM.

Compounds **1-4** (Fig. 1) were prioritized for further investigation due to their predicted affinity for a conserved glycan-binding cleft in CHI3L1, previously characterized in glycan– protein co-crystal structures.^60^ Key amino acids in CHI3L1, such as Trp99, Arg144, Val183, Asn16, Asp3, Trp212, Asp207, Arg263, Glu290, and Leu356 are located within this oligosaccharide-binding region (Fig. 2A), as characterized in PDBs 1HJV, 1HJX, 1HJW^61^. This surface-accessible pocket, which mediates interaction with short chitooligosaccharides, is both topologically stable and chemically tractable, making it a compelling target for small molecule modulation. The ability of previously reported compounds to displace chitin-or heparan sulfate-like ligands from this cleft (PDBs 8R41, 8R42, 8R4X), supports its potential as a druggable site.^60^ Based on these observations, we hypothesized that targeting the oligosaccharide-binding pocket (Fig. 2A) could alter the structural conformation or effector accessibility of CHI3L1. To explore this hypothesis, we docked reported CHI3L1 ligands (compounds **1-4**) into the binding pocket and analyzed their interaction patterns to derive a structure-based binding mode model. Additionally, the screening pharmacophores were constructed and validated based on the binding modes of compounds **2, 3**, and **4** as their interaction with CHI3L1 resulted in the highest displacement percentage of heparan sulfate.

**Fig. 1.**
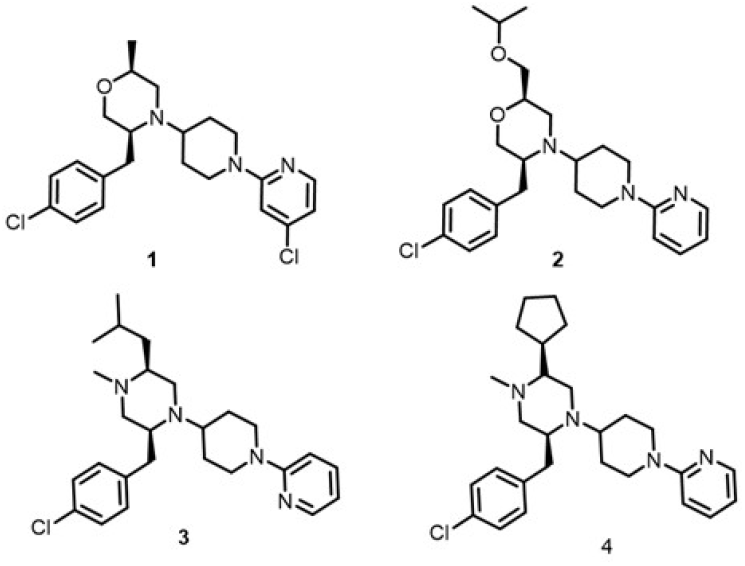
Chemical structures of compounds **1-4**, previously reported^60^ CHI3L1 ligands.

**Fig. 2.**
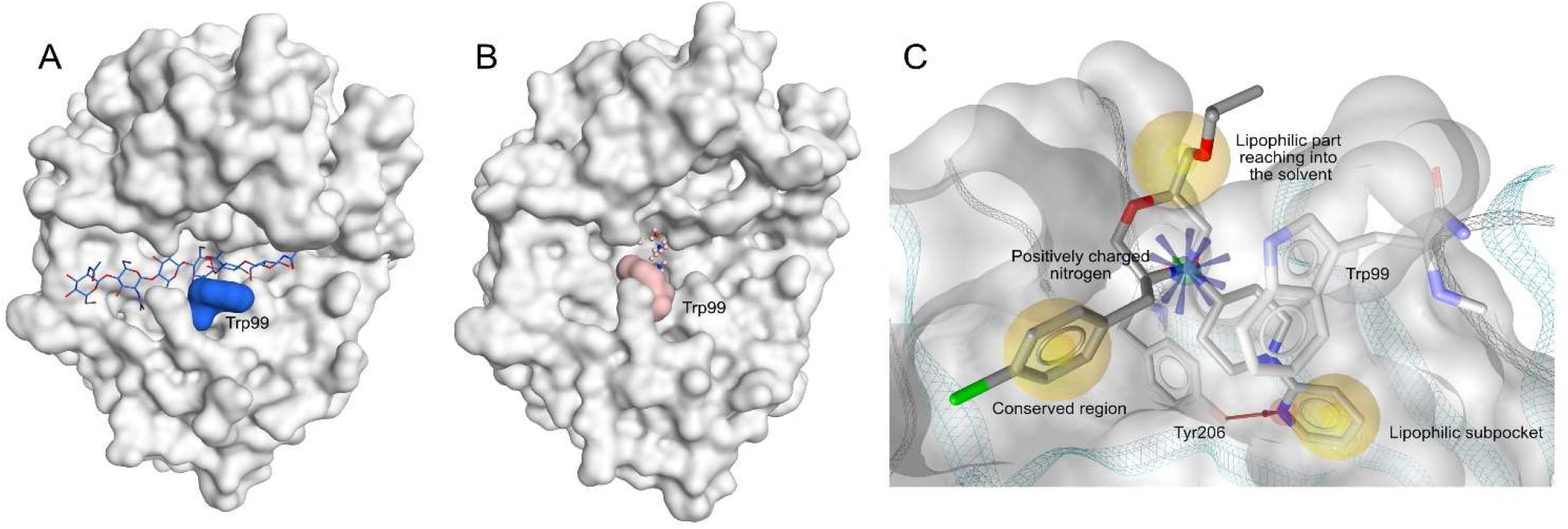
**(A)** Surface view of CHI3L1 (grey) with a oligosaccharide ligand (blue) bound in the conserved glycan-binding pocket. Trp99 (blue) is positioned at the base of the cleft; **(B)** Binding pocket of compound **1** (rose) in CHI3L1. Trp99 can adopt two different conformations, depending on the ligand class. The binding of compound **1** stabilizes Trp99 in a conformation which would clash with oligosaccharide binding; **(C)** Binding mode of compound **2** in CHI3L1. The 3D pharmacophore contains four main parts. Three of them are lipophilic features (yellow) which are situated within a lipophilic sub pocket, the conserved region of the protein and a part which reaches into the solvent. A positive ionizable feature(blue) is situated around the central nitrogen, forming hydrogen bonds and π-cation interactions to Trp99 and As9207(opposite site of Trp99, not visible).

The binding mode analysis revealed four common features which were included in the screening 3D pharmacophores. Firstly, a positively ionizable core, resulting from the charged morpholine or piperazine nitrogen. In the three co-crystallized ligands, the positively charged nitrogen formed electrostatic or hydrogen bond interactions with Asp207 and Trp99. A comparison of CHI3L1 bound to oligo-saccharides (Fig. 2A) and small molecules (e.g., compound **1** in Fig. 2B and compound **2** in Fig. 2C) showed that Trp99 can adopt two distinct binding modes depending on the ligand class. For small molecules containing the positively charged nitrogen, a π-cation interaction stabilizes Trp99 in a conformation that clashes with oligosaccharide binding. Therefore, a positive ionizable (PI) feature was included in the screening pharmacophore (Fig. 2C). Since a hydrogen bond was regularly observed between the positively charged nitrogen and Asp207, a hydrogen bond donor feature was added in one of the two main screening pharmacophores to render the PI more specific due to the restricted angle of hydrogen bonds. The analysis further revealed that all ligands align into a relatively narrow lipophilic subpocket located in the center of the oligosaccharide binding cleft (Fig. 2C). This observation is represented by a lipophilic feature in the screening 3D pharmacophore model.

Within this sub-pocket a common hydrogen bond between Tyr206 and a heteroatom of the active ligands was analyzed. As this hydrogen bond has also been recognized as a key interaction for the blocking activity of the established small compound series, a hydrogen bond acceptor feature was included as well. In one of the screening pharmacophores, an aromatic feature was included in the lipophilic subpocket, as all active ligands contain an aromatic moiety at that position, and π-π stacking was observed. As a third component of the binding mode analysis, a lipophilic contact, reaching into a conserved region of the protein, near the surface of the oligosaccharide-binding cleft was analyzed (Fig. 2C) Furthermore, compounds **2, 3** and **4** feature a lipophilic substituent at position 2 of the central morpholine or piperazine moiety, which extends into the solvent when bound to CHI3L1.

One of the two pharmacophore models used for screening (Fig. 3A) is based on compound **4**, which features a piperazine moiety as its central component, with a positively charged nitrogen atom. This pharmacophore lacks a hydrogen bond donor feature, which typically represents the hydrogen bond between the positively charged nitrogen and Asp207. The hydrogen bond acceptor feature within the lipophilic subpocket is associated with a feature for aromatic interactions. The second pharmacophore model, based on compound **2** (Fig. 3B), incorporates a morpholine moiety as the central scaffold. This pharmacophore includes a hydrogen bond donor feature for Asp207 and a hydrogen bond acceptor feature in pocket is depicted as a sphere, resulting in a less restrictive directional feature. Additionally, the lipophilic feature in the lipophilic sub-pocket, represented in vector form. However, the aromatic interaction feature is absent. ROC curve analysis of both pharmacophores revealed that model A exhibits slightly higher sensitivity, as it successfully retrieved zero false positive compounds. However, visual inspection indicated a more favorable positioning of the nitrogen in the morpholine moiety, leading to more frequent π-cation interactions with Trp99 and a hydrogen bond with Asp207.

**Fig. 3.**
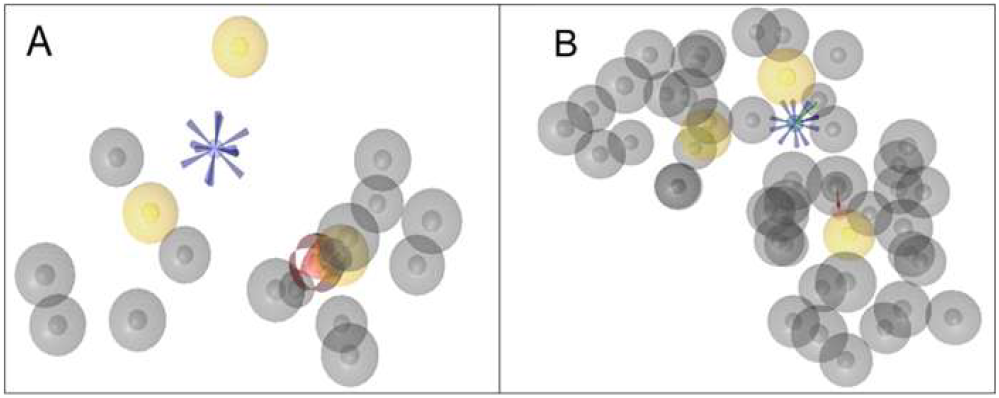
Pharmacophore models based on compounds **4 (A)** and **2 (B)**, highlighting key interaction features including hydrogen bond acceptors/donors (red and green) and hydrophobic or aromatic regions (yellow and blue) relevant to CHI3L1 binding. These two 3D Pharmacophores (**A** and **B**) were used as filters for the Enamine screening collection.

The Enamine screening collection, comprising over 4.4 million compounds, was subsequently filtered based on the constructed 3D pharmacophore models. The resulting 2071 hits were redocked into the binding site, and their conformations were rescored according to the initial screening pharmacophore. Only hits with a pharmacophore fit score above 30 were selected for further analysis. In the next filtration step, compounds with more than eight rotatable bonds or a molecular weight outside the 300–700 Da range were excluded. Finally, the remaining hits were visually inspected, focusing on the presence and geometry of the πcation interaction between the positively charged nitrogen and Trp99, as well as the hydrogen bond to Tyr206 and the two lipophilic features. The virtual screening workflow successfully identified 35 hits (Table S1) which were further evaluated for their ability for CHI3L1 binding.

Of the 35 compounds identified through virtual screening (Table S1), 33 were successfully procured from EnamineStore for experimental validation, as compounds **5** and **23** were not commercially available. To preliminarily assess their binding interactions with CHI3L1, temperature-related intensity change (TRIC) based Microscale Thermophoresis (MST) was conducted on Dianthus NT.23 Pico instrument at a single concentration of 250 μM, with each compound tested in triplicate. Compounds that showed normalized fluorescence (Fnorm) values beyond the defined threshold, calculated as the mean of the negative control ± 3 standard deviations, were selected for further evaluation. As a positive control, **G28**,^62^ a previously reported CHI3L1 binder, was included to confirm the reliability of the assay. Based on the Fnorm deviation, nine compounds were initially identified as potential hits (Fig. 4A). To further eliminate false positives due to intrinsic fluorescence or quenching effects, initial fluorescence intensities of compounds with labelled protein was compared with reference having labelled protein and DMSO (Fig. 4B). Therefore, additional autofluorescence (Fig. 4C) and quenching control assays (Fig. 4D) were performed. As a result of this filtering step, six compounds (**6, 8, 22, 31, 38**, and **39**) were identified that showed minimal interference and consistent signal stability.

**Fig. 4.**
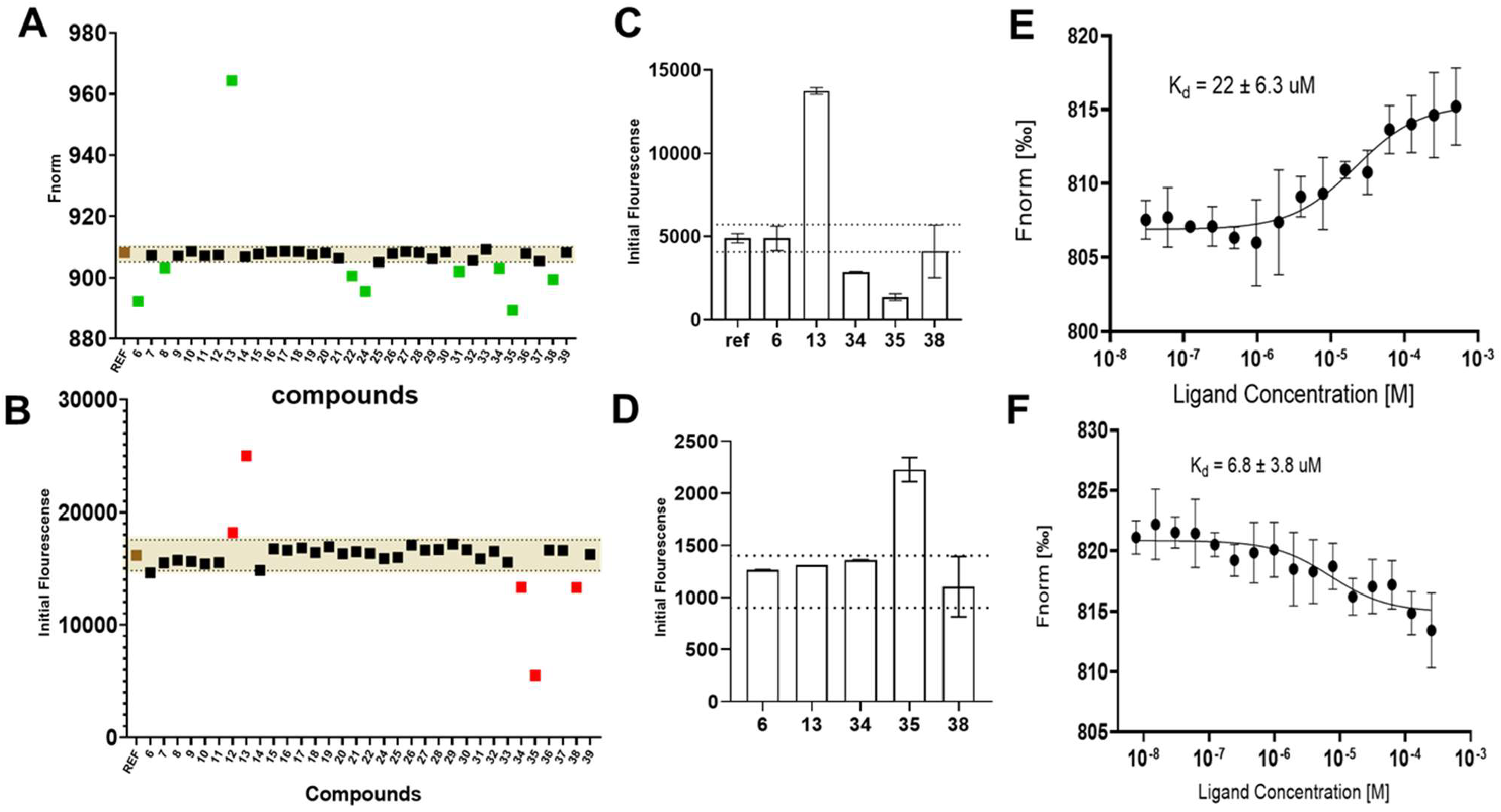
TRIC-based screening, control experiments and dose response curve of CHI3L1 binders. **(A)** Single-dose TRIC screening (250 μM, 2.5% DMSO) of Enamine library compounds with His-tagged hCHI3L1 identified potential binders. Colors indicate: blue (hits), red (autofluorescence/dye interaction), black (non-binders), orange (positive control, G28), and gray (negative control). Dashed lines represent mean negative control ± 3×SD. Fnorm values are shown as triplicate means; **(B)** Initial fluorescence of test compounds compared to the labeled protein reference; those exceeding 20% were flagged for potential interference; **(C)** Autofluorescence and **(D)** fluorescence quenching of compounds 6, 13, 34, 35, and 38 were tested to eliminate false positives due to compound interference. Assay buffer: 10 mM HEPES, 150 mM NaCl, 1% Pluronic F127, 1 mM TCEP, pH 7.4; 30 min incubation. **(E and F)** MST-based confirmation of CHI3L1 binding by compound **39** and compound **8**, respectively. The graph displays dose-dependent changes in normalized fluorescence (Fnorm [%]) plotted against increasing concentrations of each ligand.

The six hits from initial screening were then subjected to dose-response binding studies to confirm their binding affinities to CHI3L1. Of these, two compounds **8** and **39** displayed clear dose-dependent binding curves, strongly suggesting specific interactions rather than assay artifacts (Fig. 5). Compound **8** emerged as the most potent binder, exhibiting a dissociation constant (K_d_) of 6.8 ± 3.8 μM. This was followed by compound **39** with K_d_ of 22 ± 6.3 μM.

**Fig. 5.**
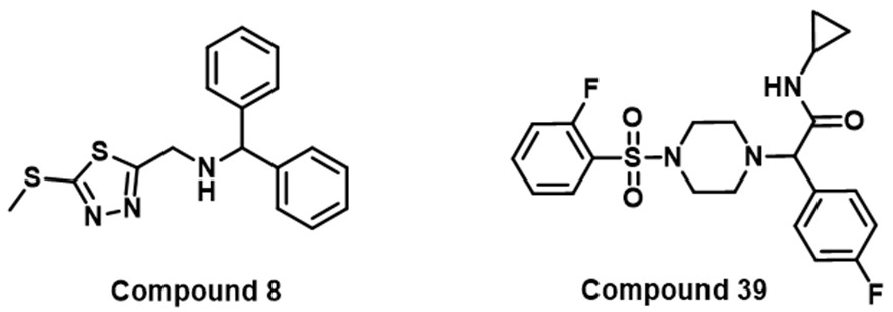
Chemical structures of the two top CHI3L1 hits (compounds **8** and **39**).

SPR was employed to validate the binding interactions of selected compounds with CHI3L1 under label-free conditions. As a highly sensitive technique that enables realtime monitoring of molecular interactions, SPR serves as a valuable complement to microscale thermophoresis (MST). To corroborate the MST findings, we assessed the binding affinities of compounds **8** and **39** using SPR. Compound **8** showed a clear, dose-dependent binding response, yielding a dissociation constant (K_d_) of 5.69 μM. Similarly, compound **39** demonstrated measurable binding with a K_d_ of 17.09 μM (Fig. 6).

**Fig. 6.**
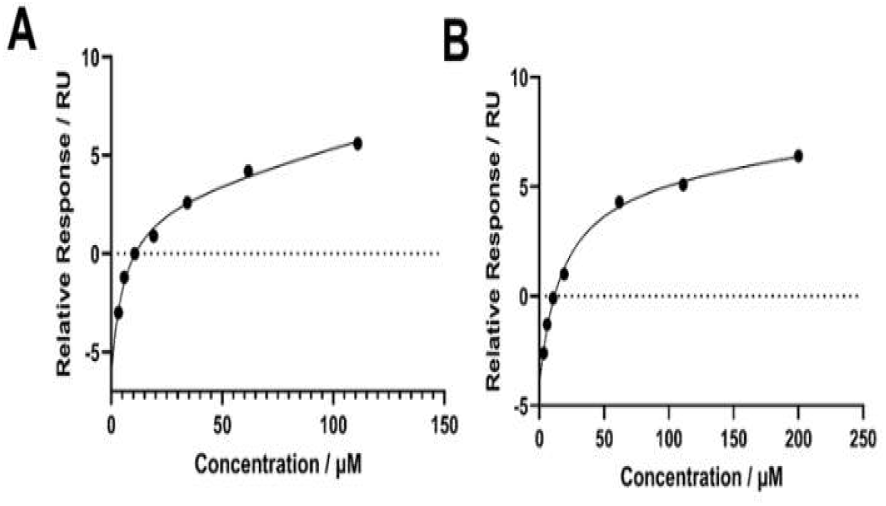
SPR-based binding affinity analysis of compounds 8 and 39 towards hCHI3L1. Dose-dependent binding curves of **(A)** compound 8 and **(B)** compound **39** to hCHI3L1 as measured by SPR. His-tagged hCHI3L1 protein (8.0 μg/mL) was immobilized on a CM5 sensor chip, and serial concentrations of each compound (0–250 μM) were injected in running buffer using a single-cycle kinetic model.

To further support the observed CHI3L1 binding affinity of the hit compounds, a detailed docking analysis was performed for compounds **8** and **39** within the active site of hCHI3L1 (Fig. 7). The docking results revealed that both compounds stabilize the side chain of Trp99 in a conformation that is unfavorable for oligosaccharide binding. This stabilization is mediated through charged nitrogen atom of the central amine in compound **8** or the piperazine ring in compound **39**. Additionally, hydrogen bonding interactions further support the binding affinity of these ligands: compound **8** forms a hydrogen bond between the 1,3,4-thiadiazole ring and Tyr206, whereas compound **39** engages Tyr206 through its sulfonamide group. Furthermore, both compounds exhibit a three-part lipophilic architecture, with distinct segments extending into the lipophilic sub-pocket, the conserved region of the protein, and the solvent-exposed area. These findings provide structural insights into the molecular basis of ligand binding and offer a rationale for the inhibitory potential of the identified compounds.

**Fig. 7.**
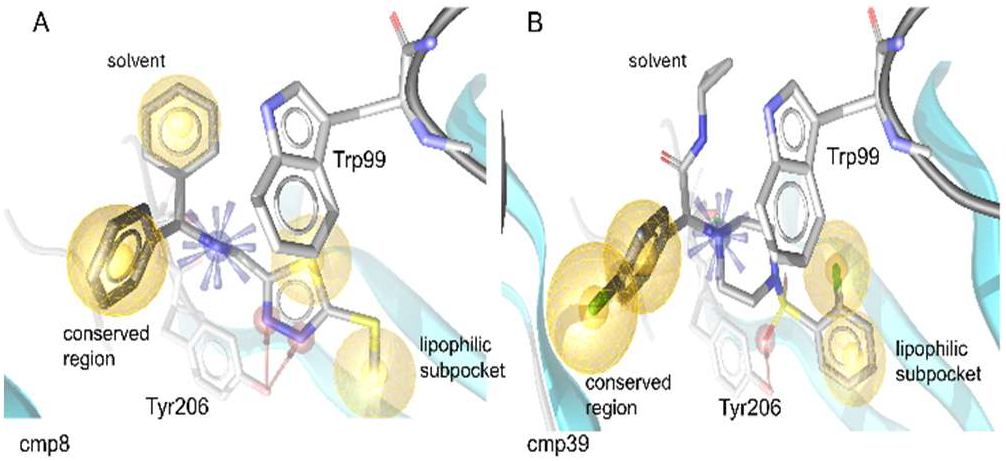
Docking poses of compounds 8 and 39 in hCHI3L1. Both compounds stabilize Trp99 in the position which is unfavorable for oligosaccharide binding by forming a π-cation interaction to the positively charged nitrogen of the central amine in **8** or piperazine moiety in **39**. The binding modes are stabilized by hydrogen bonds between Tyr206 and the 1,3,4-thiadiazole moiety in compound **8** and the sulfonamide moiety in compound **39**. The three lipophilic parts reach into the lipophilic subpocket, the conserved region of the protein and the solvent.

To evaluate functional activity in a physiologically relevant system, we tested compounds **8** and **39** in a multicellular 3D GBM spheroid model^62^ comprising GBM, endothelial, and macrophage cells. This model was selected as it more accurately mimics the complex tumor microenvironment of GBM. HB, a known CHI3L1–STAT3 interaction disruptor,^51^ was used as a control. Treatment with compound **8** resulted in a clear, dose-dependent reduction in spheroid viability, with significant effects at 10 μM (Fig. 8A). In contrast, compound **39** exhibited only modest activity, with viability reductions reaching significance only at 25 μM and higher (Fig. 8A). These outcomes are consistent with their respective CHI3L1 binding affinities, as compound **8** demonstrated ~3-fold stronger CHI3L1 binding than compound **39**.

**Fig. 8.**
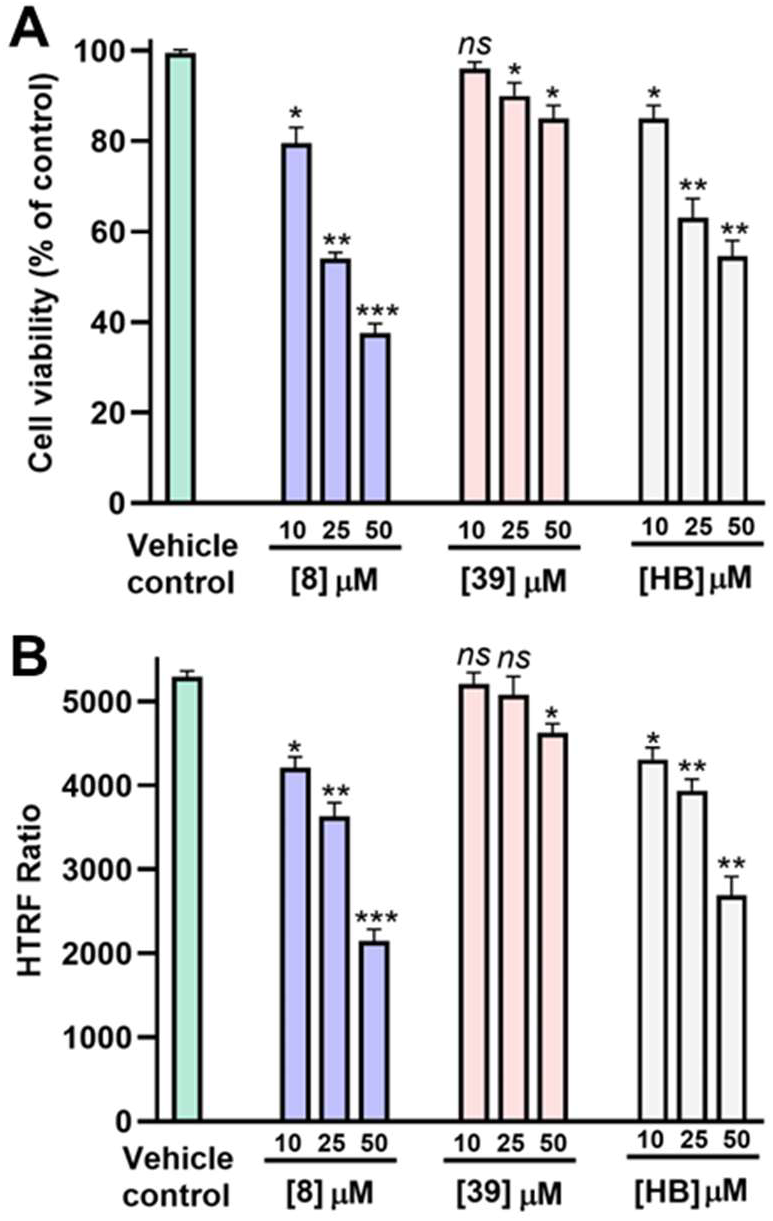
Evaluation of the therapeutic potential of compounds 8 and 39 in GBM spheroids. **(A)** Cell viability of GBM spheroids (as % of untreated control) upon incubation with increasing concentrations of the tested compounds (**8, 39**, and HB) after 72 h incubation. **(B)** Reduction in phospho-STAT3 levels, as determined by HTRF phospho-STAT3 kit from Revvity (Cat# 62AT3PET), in GBM spheroids upon incubation with the tested compounds (**8, 39**, and HB) after 72 h incubation. * *p* < 0.05, ** *p* < 0.01, *** *p* < 0.001, and (*ns*) denotes nonsignificant relative to vehicle control. Data are representative of three independent experiments.

We next measured phospho-STAT3 levels using a homogeneous time-resolved fluorescence (HTRF) assay. Compound **8** showed dose-dependent inhibition of pSTAT3, mirroring the effects of HB, a known CHI3L1–STAT3 disruptor (Fig. 8B). Compound **39** had minimal impact on pSTAT3 except at 50 μM. Together, these results highlight compound **8** as a potent CHI3L1 pathway inhibitor, capable of reducing GBM spheroid viability and suppressing STAT3 activation in a physiologically relevant 3D model.

Overall, using a pharmacophore-based virtual screening strategy, we identified two small molecules with micromolar binding affinity to CHI3L1, as confirmed by MST and SPR. These compounds represent promising CHI3L1-binding scaffolds with validated biophysical engagement, offering a foundational advance in targeting this challenging, non-enzymatic immunomodulatory protein. While further investigation is needed to elucidate the full biological consequences of CHI3L1 inhibition, initial studies indicate that compound **8** not only binds CHI3L1 but also reduces spheroid viability and downregulates STAT3 phosphorylation, hallmarks of disrupted CHI3L1 signaling. These findings suggest potential for therapeutic modulation of CHI3L1 in diseases such as GBM, where it contributes to immune suppression and tumor progression. Beyond these specific findings, this work exemplifies how rational in silico design, anchored in structural pharmacophore modeling and reinforced by orthogonal biophysical validation, can yield tractable small molecule ligands for proteins historically considered undruggable. Moving forward, these hits provide a promising platform for medicinal chemistry optimization and mechanistic dissection of the role of CHI3L1 in cancer and inflammation, ultimately enabling the development of targeted modulators to probe and manipulate this critical signaling axis.

## Supporting Information

The Supporting Information is available free of charge on the ACS Publications website.

Computational methods, biophysical screening assays, cellular evaluation, and SMILES of selected compounds (PDF)

## Author information

### Author Contributions

The manuscript was written through contributions of all authors. All authors have given approval to the final version of the manuscript.

## Abbreviations

CD44: Cluster of Differentiation 44
CHI3L1: Chitinase-3-like protein 1
Gal3: Galectin-3
Gal3BP: Galectin-3 Binding Protein
GBM: glioblastoma
HB: hygromycin B
HTRF: homogeneous time-resolved fluorescence
IL-13Rα2: IL-13 receptor alpha 2
KD: equilibrium dissociation constant
MST: microscale thermophoresis
PMT: proneural-to-mesenchymal transition
PPI: protein–protein interactions
RAGE: receptor for advanced glycation end-products
SPR: surface plasmon resonance
STAT3: signal transducer and activator of transcription 3.

## Notes

The authors declare no competing financial interests.

## Acknowledgments

We gratefully acknowledge financial support from the National Institute of Neurological Disorders and Stroke under grant number R01NS136524 (PI: Gabr)

