## Supporting Information for "Virtual Screening–Guided Discovery of Small Molecule CHI3L1 Inhibitors with Functional Activity in Glioblastoma Spheroids"

**Contents**

| 1. | Computational Methods | S2 |
| --- | --- | --- |
| 2. | Biophysical Screening Assays | S3 |
| 3. | Screening in GBM spheroids | S4 |
| 4. | SMILES of selected compounds | S5 |
| 5. | References | S6 |

**1. Computational Methods**

1.1. **Retrospective SAR analysis of known active ligands**

Reported active small molecule ligands were built and protonated in LigandScout (Inte:Ligand, v.4.43, Vienna Austria) and MOE (Molecular Operating Environment, v.2022.02 Chemical Computing Group ULC, 910-1010 Sherbrooke St. W., Montreal, QC H3A 2R7 ,2025.), stereochemistry was visually inspected. The crystal structures of 8R41, 8R42 8R4X^1^ were downloaded from [www.rcsb.org](http://www.rcsb.org) and manually prepared and protonated in MOE. Ligands were then docked into the binding site of compound **1**^1^ in 8R4X to reproduce the reported SAR. 10 docked poses were generated in Gold for each ligand, poses which didn’t enter the binding pocket were discarded. All co-crystallized ligand conformations were found in the docking, poses can be regarded as validated. For the construction of the 3D screening pharmacophores in LigandScout, all docked poses of the reported active ligands were visually inspected to generate a basic binding mode hypothesis which is described in detail in the result part.

1.2. **Prospective virtual screening**

The Enamine Screening Collection was downloaded and prepared using a Knime workflow. 4 3D pharmacophores based on compounds **2**, **3** and **4** were validated using receiver operating curves (ROC) as part of the iscreen program in LigandScout. The reported actives were classified as actives and a set of Dud-e decoys was classified as inactives. (generated on <https://dude.docking.org/> based on reported actives). Two out of four 3D pharmacophores resulted in promising ROC curves.

Virtual screening rounds were then performed using iscreen on the Computing Cluster of the Molecular Design Lab, Freie Universität Berlin. Hits were redocked into the binding site using Gold and their conformations were rescored according to the initial screening pharmacophores. Only hits with a pharmacophore fit score above 30 were selected for further analysis. In the next filtration step, compounds with more than eight rotatable bonds or a molecular weight outside the 300–700 Da range were excluded using LigandScout. Finally, the remaining hits were visually inspected, and the final hits were selected considering pharmacophore feature occurrence, binding pocket shape and chemical plausibility.

2. **Biophysical Screening Assays:**

**2.1. Microscale Thermophoresis (MST)**

**2.1.1. Initial Screening via Single-Point MST**

The His-tagged CHI3L1 protein (Cat. #CH1-H5228, Acro Biosystems, Newark, DE, USA) was fluorescently labeled using the second-generation RED-tris-NTA dye from the Monolith His-Tag Labeling Kit (Cat. #MO-L018, NanoTemper Technologies, München, Germany) as per the supplier’s guidelines. For each experiment, a 100 nM dye solution was incubated with 200 nM of the target protein in PBS buffer having 0.05% tween 20 at ambient temperature, shielded from light, for 30 minutes.

Post-labeling, the protein was diluted in assay buffer (10 mM HEPES, 150 mM NaCl, 0.1% Pluronic F-127, 1 mM TCEP, 8% DMSO, pH 7.4) and mixed with the test compound to obtain final concentrations of 20 nM of protein) and 250 μM of compound with 4% DMSO. The mixture was allowed to incubate for 30 minutes at room temperature in the dark, followed by brief centrifugation (1000 × g for 30 seconds) and transferred into the Dianthus NT.23 Pico (NanoTemper Technologies). A control sample containing only DMSO in assay buffer was included in all runs. All experiments were conducted in triplicate, and mean values were reported. To assess intrinsic fluorescence from test compounds, each compound was diluted in assay buffer from a DMSO stock to a concentration of 250 μM. The samples were kept in the dark at room temperature for 30 minutes, then centrifuged at 1000 × g for 30 seconds. The Dianthus NT.23 Pico was used to measure fluorescence. Control assays contained buffer with DMSO only. Each assay was performed in technical triplicates, and results were averaged and reported as mean ± SD. A quenching control experiment was performed by mixing each test compound (final concentration: 250 μM) with RED-tris-NTA 2nd generation dye (10 nM) in assay buffer. The mixture was incubated for 30 minutes at room temperature, protected from light, and centrifuged at 1000 × g for 30 seconds. Fluorescence readings were acquired using the Dianthus NT.23 Pico. Control samples consisted of dye with DMSO in buffer. Measurements were completed in triplicate, with results expressed as mean ± standard deviation.

**2.1.2. Dose dependent MST Binding Analysis**

Dose–response MST experiments were carried out on hit compounds, using the Monolith NT.115 instrument (NanoTemper Technologies). CHI3L1-His protein (Cat. #CH1-H5228, Acro Biosystems) was labeled with the RED-tris-NTA dye (Cat. #MO-L018) in HEPES buffer. Each compound was prepared as a 16-point serial dilution starting from 1 mM down to low nanomolar concentrations in PBST buffer, yielding a final DMSO concentration of 4% upon mixing with protein. After combining labeled protein with each dilution at a 1:1 ratio, the solutions were incubated for 30 minutes, centrifuged briefly, and then loaded into MST capillaries. MST readings were taken using 60–80% LED intensity in the red channel and medium to high IR-laser power. Data interpretation was performed with MO.Affinity Analysis v2.3 (NanoTemper Technologies).

**2.2. Surface Plasmon Resonance (SPR) Binding Assay**

SPR was employed to assess the interaction between compound G28 and CHI3L1-His protein using the Biacore 8K system (Cytiva). A single-cycle kinetic method was applied, wherein increasing concentrations of G28 were injected sequentially over a sensor chip with immobilized protein. All measurements were performed at 25 °C. Sensorgrams were corrected by subtracting background signals obtained from a reference flow cell and buffer-only injections. Data evaluation was carried out using Biacore™ Insight Evaluation Software (Cytiva).

**3. Screening in GBM spheroids**

The GBM spheroids were prepared as previously described.^2^ U-87 MG GBM cells (ATCC, Cat# HTB-14) were cultured in DMEM (ATCC, Cat#30-2002) containing 4.5 g/L glucose and 2 mM L-glutamine, supplemented with streptomycin and penicillin. HMEC-1 (ATCC) were maintained in MCDB131 medium with 10 mM L-glutamine, 10 ng/ml FGF, and 1 µg/ml hydrocortisone. Briefly, U-87 MG and HMEC-1 cells were co-seeded with macrophages on low-adhesion 96-well plates at 2 × 10³ cells per well in 100 µl of the tested compounds at varying concentrations (10, 25, and 50 µM) or control media. After 72 hours, cell viability was assessed using the CCK-8 assay (MedChemExpress, Cat# HY-K0301) according to the manufacturer’s recommended protocol. Absorbance at 450 nm was measured using a microplate reader.

Assessment of phospho-STAT3 levels was determined by HTRF phospho-STAT3 kit from Revvity (Cat# 62AT3PET) using the manufacturer’s recommended protocol. All experiments were conducted in triplicate.

**Table S1. Smiles of selected hits from Enamine library as potential CHI3L1 hit compounds.**

| Manuscript ID | Catalog ID | Smiles |
| --- | --- | --- |
| 5 | Z2826283839 | [N+H2](C[C@H](N1Cc2c(scc2)CC1)c1ccccc1)C1CC(n2nccc2)C1 |
| 6 | Z125428276 | [N+H](Cc1occc1)(Cc1ccc(-n2nccc2)cc1)Cc1ccccc1 |
| 7 | Z989733528 | O=C(NC1CC1)[C@@H]([N+H]1CCC(c2[nH]ncc2)CC1)c1ccccc1 |
| 8 | Z826980060 | S(C)c1sc(C[N+H2]C(c2ccccc2)c2ccccc2)nn1 |
| 9 | Z195570878 | O=C(Nc1noc(C)c1)[C@@H](CC)[N+H]1CCN(c2ncccc2)CC1 |
| 10 | Z3998746959 | FC(F)(CN[C@H](C(=O)NC1CC1)c1ccccc1)c1ncccc1 |
| 11 | Z1551102255 | Brc1cc2[C@H]([N+H2]Cc3[nH]nc(CC)c3)[C@@H](C)Cc2cc1 |
| 12 | Z840305718 | C([N+H]1C[C@@H](C)O[C@H](c2ccccc2)C1)c1nc2n(C(C)=CC(C)=N2)c1 |
| 13 | Z1185943433 | O=C(Nc1scnn1)C[N+H]([C@@H](c1ncccc1)c1ccccc1)C |
| 14 | Z1142756634 | O=C(NC(C)C)[C@H]([N+H]1Cc2n(c(C3CC3)nn2)CC1)c1ccccc1 |
| 15 | Z1603789773 | O=C(OCC)N1CCC([N+H2][C@@H](c2cnccc2)c2ccccc2)CC1 |
| 16 | Z642877720 | O[C@H](C[N+H]1[C@@H](c2ccccc2)c2c(scc2)CC1)Cn1nccc1 |
| 17 | Z991108824 | O=C(NC1CC1)[C@H]([N+H]1C[C@@H](c2sccn2)OCC1)c1ccccc1 |
| 18 | Z1438282549 | Brc1ccc([C@H]([N+H2]Cc2ncc(C)nc2)C2CCC2)cc1 |
| 19 | Z224853528 | Fc1ccc(NC(=O)[C@@H](CC)[N+H]2CCN(c3sccn3)CC2)cc1 |
| 20 | Z1620872904 | FC(F)(F)CN1C(=O)C[C@@H]([N+H2][C@H](c2ncccc2)c2ccccc2)C1 |
| 21 | Z3634835945 | Fc1ccc([C@@H]([N+H2]C2CC(F)(C[N+H3])C2)c2cnc3c(c2)cccc3)cc1 |
| 22 | Z1130120284 | Fc1ccc([C@H]([N+H2]Cc2oc(C3CC3)nn2)C2CCC2)cc1 |
| 23 | Z3359948765 | [N+H2]([C@H](CC)c1sccn1)C1CCN(c2nnccc2)CC1 |
| 24 | Z2058143891 | C([N+H]1[C@@H]([C@@H](C)OCC1)c1onc(C2CC2)n1)c1nc(C)on1 |
| 25 | Z241896688 | O=C(Nc1onc(C)c1)C[N+H]1[C@H](CC)c2c(scc2)CC1 |
| 26 | Z1731222549 | Fc1c(N2C[C@@H]([N+H2][C@H](CC)c3c(O)cccc3)CC2)nccc1 |
| 27 | Z2610539188 | Fc1cc2OC(C)(C)C[C@@H]([N+H2]C[C@@H](O)c3ncccc3)c2cc1 |
| 28 | Z1567594442 | C(C)c1c(C[N+H]2CCN(c3nnccc3)CC2)c2c(o1)cccc2 |
| 29 | Z1176669137 | Fc1ccc(NC(=O)[C@H]([N+H]2CCC([C@@H](O)C)CC2)c2ccccc2)cc1 |
| 30 | Z100502520 | O=C(C[C@@H]1C=CCC1)N1CC[N+H](C(c2ccccc2)c2ccccc2)CC1 |
| 31 | Z986034082 | O=C(Nc1ccccc1)[C@H]([N+H]1CCC(c2nc(C)on2)CC1)c1ccccc1 |
| 32 | Z32558286 | O=C(NC[C@@H]([N+H]1CCC(C)CC1)c1sccc1)[C@H]1Oc2c(OC1)cccc2 |
| 33 | Z2442140468 | Fc1ccc(C([N+H]2CCN(C(=O)[C@@H](C(=O)[O-])C)CC2)c2ccc(F)cc2)cc1 |
| 34 | Z2617090746 | O=C1n2nccc2NC(C[N+H]2CC(C)(C)Cc3n(-c4ccncc4)ncc3C2)=C1 |
| 35 | Z56174695 | Oc1c([C@@H]([N+H]2CCN(c3ncccc3)CC2)c2sccc2)ccc2c1nccc2 |
| 36 | Z118229966 | S(CC[C@H]1C(=O)N(C[N+H]2[C@@H](c3sccc3)c3c(scc3)CC2)C(=O)N1)C |
| 37 | Z238821776 | Fc1ccc(NC(=O)[C@H]([N+H2]C2CCN(C(=O)c3occc3)CC2)c2ccccc2)cc1 |
| 38 | Z44521666 | O=[N+]([O-])c1c(NC(=O)[C@H]([N+H]2CCN(c3ncccn3)CC2)c2ccccc2)ccc(C)c1 |
| 39 | Z224260766 | S(=O)(=O)(N1CC[N+H]([C@H](C(=O)NC2CC2)c2ccc(F)cc2)CC1)c1c(F)cccc1 |
